# Transparency and open science reporting guidelines in sleep research and chronobiology journals

**DOI:** 10.1101/2020.06.26.172940

**Authors:** Manuel Spitschan, Marlene H. Schmidt, Christine Blume

**Affiliations:** Department of Experimental Psychology, University of Oxford, United Kingdom; Centre for Chronobiology, Psychiatric Hospital of the University of Basel (UPK), Switzerland; Transfaculty Research Platform Molecular and Cognitive Neurosciences, University of Basel, Switzerland

## Abstract

“Open science” is an umbrella term describing various aspects of transparent and open science principles. The adoption of open science principles at different levels of the scientific process (e.g., individual researchers, laboratories, institutions) has been rapidly changing the scientific research landscape in the past years, but uptake of these principles differ from discipline to discipline. Here, we asked to what extent journals in the field of sleep and chronobiology research encourage or even require following transparent and open science principles in their author guidelines. To this end, we scored the author guidelines of a comprehensive set of 28 sleep and chronobiology journals, including the major outlets in the field, using the standardised Transparency and Openness (TOP) Factor. This instrument rates the extent to which journals encourage or require following various aspects of open science, including data citation, data transparency, analysis code transparency, materials transparency, design and analysis guidelines, study pre-registration, analysis plan pre-registration, replication, registered reports, and the use of open science badges. Across the 28 journals, we find low values on the TOP Factor (median [25^th^, 75^th^ percentile] 2.5 [1, 3], min. 0, max. 9, out of a total possible score of 28). This suggests an opportunity for sleep and chronobiology journals to further support the recent developments by implementing transparency and openness principles in their guidelines and making adherence to them mandatory.

## Introduction

During the last years, the open science movement gained increasing popularity and is rapidly changing the way science is done, especially among early career researchers striving to improve scientific practice and overcome deficits in the current scientific *status quo*^*1,2*^. The term “open science” is relatively ill-defined and includes a range of different methods, tools, platforms, and practices that are geared to improving the quality of science through transparency^3^. At present, it is still largely up to individual researchers and research groups to decide to what extent they want to engage in open science practices and incentives that may promote open science are rare. Journals as the main outlets for archival scientific dissemination can support the movement and offer ways to make the scientific process more open, reproducible, and emphasise good scientific practice. They may even speed up the process by requiring authors to adhere to open science standards. However, to what extent do journals in the fields of sleep and chronobiology encourage or even require following the standards of open science?

The scientific fields of sleep research and chronobiology concern all aspects of sleep and circadian rhythmicity. As almost all aspects of physiology and behaviour are under some type of circadian control, this cluster of scientific fields is fundamentally interdisciplinary, employing a wide variety of methodologies. Therefore, this research area is very heterogeneous, drawing from different ‘core’ disciplines (including neuroscience, psychology, molecular biology, and others), each with their own scientific history, and the degree to which open science principles are adopted may vary widely.

In this study, we asked to what extent scientific journals specialised on sleep research and chronobiology lay out open-science principles in their author guidelines. Inspired by previous publications in other fields^4,5^, we assessed the implementation of research transparency and openness in journal guidelines using the quantitative Transparency and Openness Factor^6^ (TOP Factor; https://topfactor.org/). The TOP Factor contains ten sub-scales, corresponding to different aspects of openness and transparency in scientific research, reflecting the Transparency and Openness Promotion Guidelines^7^: data citation, data transparency, analysis code transparency, materials transparency, design and analysis guidelines, study pre-registration, analysis plan pre-registration, replication, registered reports, and open science badges.

The TOP Factor recognises different levels relating to mentioning, encouraging, requiring and enforcing specific transparency and openness practices, which are implemented in a verbally anchored rating scheme. *Data citation* refers to the citation of data in a repository using standard means, including a digital object identifier (DOI). *Data, analysis code*, and *materials transparency* refers to making data, analysis code and materials available as part of the journal submission. The category *Design and analysis guidelines* refers to the inclusion of instruments describing the study design and analysis formally, such as the Preferred Reporting Items for Systematic Reviews and Meta-Analyses (PRISMA) or Consolidated Standards of Reporting Trials (CONSORT) standards. *Study pre-registration* and *analysis pre-registration* refers to the pre-registration of data collection and/or analysis prior to their excitation. *Replication* refers to an explicit desire of the journal to include articles not based on novelty. The category *Registered reports* refers to prospective peer review, i.e. evaluation of a manuscript submitted to a journal prior to data collection and/or data analysis. Registered reports have recently gained significant traction, with a few high-profile journals, including Nature Human Behaviour and PLOS Biology, accepting this ‘frontloaded’ article format. *Open-science badges* refers to the use of so-called badges, which are awarded if a paper adheres to specific standards, thereby providing an incentive for promoting transparency and openness^8^. In summary, the TOP factor covers major dimensions of open science and provides a helpful and standardised tool that allows to compare between journals or fields the extent to which they encourage or require adherence to open science principles.

## Methods

### TOP Factor

The TOP Factor (Transparency and Openness Factor) is a quantitative score summarising the presence, requirement, and enforcement of transparent and open science practices in journals. It includes a total of ten sub-scales, of which nine score 0-3, and one scores 0-2, thereby resulting in a maximal summed score of 29. Higher values indicate a higher degree of adherence to the TOP practices.

### Journal identification

Journals to be included in the rating were identified using a hybrid pre-registered strategy (https://doi.org/10.17605/OSF.IO/QNSBM):

- *Primary strategy*. Relevant journals were identified using search on the Web of Science Master Journal List (WoS MJL; https://mjl.clarivate.com/). The search terms, entered in separate searches, were:
  - “sleep”
  - “chronobiology”
  - “circadian”
  - “biological rhythms”
  - “dream”
- The search results were merged, and duplicates were removed. We validated our search strategy by confirming that all journals listed in a recent publication on sleep research journals^9^ were identified using this strategy.
- *Secondary strategies*. In addition to the primary search strategy, we used two supportive secondary strategies to identify relevant journals that may have been missed in the primary strategy:
  - Own domain-relevant expertise in sleep and chronobiology;
  - Informal consultation with a senior researcher with >25 years of experience in the field.
- *Validation*. We validated our search strategy by confirming that the above search terms produce the same list of journals in MEDLINE (https://www.ncbi.nlm.nih.gov/nlmcatalog?term=currentlyindexed%5BAll%20Fields%5D%20AND%20currentlyindexedelectronic%5BAll%20Fields%5D&cmd=DetailsSearch).

In addition to this strategy, we found two additional journals via the search for TOP signatories, and one through a search in the National Library of Medicine (NLM; https://www.nlm.nih.gov/).

### Journal meta-data extraction

We extracted the 2018 Journal Impact Factor (JIF) and the 5-year Journal Impact Factor from the *Clarivate Analytics InCites* platform. The 2018 JIF was available for 15 out of 28 journals (53.6%), and the 5-year JIF was available for all of these 15 except one (14 out of 28; 50%). We obtained the NLM ID using search on the NLM data base, from which we also extracted the MEDLINE indexing status and the first year of publication. Information regarding support by scientific or professional societies (11 out of 28, 39.2% of journals were not at present supported by a society) was extracted from both the NLM entry, and the journal website. Three journals accepted submissions in a language other than English.

### Journal guidelines extraction

We consulted the journal websites for author guidelines. Where possible, we archived journal guidelines either locally, or on the Internet Wayback Machine (https://archive.org/web/). One journal, *Sleep Medicine Reviews*, did not have any public author guidelines available, as it is an invite-only journal.

### Scoring and conflict resolution

Three scorers (authors of this study, M.S., M.H.S, and C.B.) independently assessed the 28 identified journals’ TOP Factors in a total of 280 individual ratings (28 journals × 10 rating categories). In a first pass, the three scorers agreed in 75% of all ratings (210 out of 280 ratings). We then discussed and resolved major sources of discrepancy (e.g., we agreed that a clinical trial registration counted as preregistration), resolved some per-item disagreements and rescored the categories “Data citation” (initial disagreement rate: 13/28), “Reporting guidelines” (initial disagreement rate: 13/28) and “Study pre-registration” (initial disagreement rate: 19/28, see above) independently in a second pass. At the end of this second pass, all ratings agreed. All scorings were completed between mid-May and mid-June 2020.

### Data and materials availability

The scoring book including intermediate scoring stages is included in the supplementary material (Supplementary Material S1). The TOP Factor scoring rubric we used is likewise included (Supplementary Material S2).

## Results

### Low explicit implementation of transparency and openness in sleep research and chronobiology journals

Across the 28 journals we examined, we find a total median TOP Factor of 2.5 (25^th^ percentile 1, 75^th^ percentile 4, minimum 0, maximum 9, IQR 3) out of a maximum of 29 points. The three journals scoring highest on the TOP were *Clocks & Sleep* (9), *Sleep Science and Practice* (7), and *Sleep and Vigilance* (6). Interestingly, these three journals were founded no earlier than 2017. Our results compare to the low uptake of transparency and openness principles in the recent original and cross-sectional follow-up studies investigating transparency and openness in pain research^4,5^. Across ten journals in the pain research field, a median TOP Factor of 3.5 (IQR 2.8) was found. We see the low transparency and openness scores in sleep research and chronobiology journals as an opportunity to revisit how we do science, and how we report it.

### Lack of a standard specification for journal guidelines

Across the 28 journals we examined, author guidelines were widely varying in their accuracy, detail, and organisation of information. Many journals appeared to follow standard publisher guidelines, with very little or no modifications for the specific journal and often even referred to the publisher guidelines for further information. An additional challenge comes from the fact that the public-facing journal guidelines are not fully indicative of the process that the journal will implement, as further guidelines or requirements may be hidden in the submission system, or in correspondence with the journal during peer review or after acceptance of the article. For example, it is unclear to what extent a rule will be enforced in the submission process when the guidelines say that authors ‘will be asked to’ do something. Fundamentally, this unseen information may limit the extent to which public author guidelines are truly reflective of the enforcement of transparency and openness principles in a given journal. In one instance, the editorial celebrating the inaugural issue of the journal stated that its welcomes Registered Reports, but at present, the author guidelines do not explicitly state this^10^. Unless one was to consult this additional information, it would remain unknown. One way to improve transparency and openness may be to devise a standard specification schema for submission guidelines, reflecting the categories in the TOP Factor.

## Discussion

### Ambiguity in transparency and openness standards

There can be large ambiguity in the extent that a journal implements specific transparency and openness standards. Take, for example, the category “Study pre-registration”. There are four levels in this category: Level 0: *Journal says nothing*; Level 1: *Articles will state if work was preregistered*; Level 2: *Article states whether work was preregistered and, if so, journal verifies adherence to preregistered plan*; Level 3: *Journal requires that confirmatory or inferential research must be preregistered*. According to the TOP Guidelines (v1.0.1), Level 1 is satisfied if the research was registered in an independent, institutional registry, specifying “study design, variables, and treatment conditions prior to conducting the research”, leaving the level of detail open and rendering scorings ambiguous. And indeed, there is a debate and confusion regarding the use of the terms *registration* vs. *pre-registration*^*11*^. While the registration of a clinical trial in a trial registry can be relatively lightweight, containing only minimal details, a pre-registration (as used in the open science community) typically refers to the prospective specification of concrete study details, including methodology, sample size, and analysis plan prior to data collection^12^. In more detail, the registration of a clinical trial in a registry such as clinicaltrials.gov on the one hand, and the pre-registration of analysis procedures and hypotheses prior to conducting the research on the other hand, mostly serve fundamentally different purposes, which is reflected in their nature too. First, clinical studies, which have not been registered, are impossible to publish in respected journals rendering the process a necessity rather than a self-imposed step to improve scientific transparency. Furthermore, if authors register a clinical trial for instance on the German Clinical Trials Register, they have to provide a short description of the trial, name the study goals, describe the intervention, name the primary endpoints, inclusion and exclusion criteria, the final sample size (without rationale), and the sponsor. Clearly, although the degree of detail is of course also subject to variation among pre-registered studies, the required level of detail for registering clinical studies is extremely low with accountability consequently likewise being very low. In some legislations (such as Switzerland), the submission of ethics application as a clinical trial (which is required for some studies that modify sleep schedules), even automatically deposits the study in the (Swiss) clinical trial registry. Further developments of the TOP guidelines should therefore reflect the extent to which something has been preregistered, possibly also including at which time point during the scientific process the registration has taken place. Likewise, journals should be clear about what level of preregistration they expect.

### Linguistic details

*When is ‘should’ mandatory?* The author guidelines also differed in the degree they used language to specify requirements. For example, many journals “encouraged” authors to do something, but the use of this term basically carries no power – you may also just ignore it. The use of the verb “should” may be intended to signal mandatory requirements, but it leaves the possibility of ignoring the requirement. Likewise, journals that “ask authors to do something” may still allow exceptions. This may not only be favourable for authors, who do not comply with the requirements, but also allows editors to treat some submissions different from others. Moving forward, journals should state what aspects specified in guidelines are recommendations, what are requirements, and what the consequences for not meeting requirements are. To promote open science culture, it is clear that ‘hard’ requirements need to replace ‘soft’ encouragement. This is because pre-registering is an additional step, it costs time and many researchers are still not convinced it will eventually pay off. If, in addition to this, the reward is too low or non-existent, or there are no tangible negative consequences, even diligent scientists become a bit lazy.

### Open review as an additional open science dimension

Some journals, including eLife, PLOS, and Clocks & Sleep, now offer posting of the pre-publication peer-review, with the possibility of naming the reviewers (if they agree). This does not only make the journey of an article from submission to publication transparent. It also curtails unreasonable requests during peer review and may encourage reviewers to provide constructive feedback oriented towards the best scientific outcome. We therefore encourage to include “open review” as an additional category in future developments of the TOP guidelines.

## Conclusion

In a comprehensive analysis of the author guidelines for 28 sleep and chronobiology journals, we have found low evidence for explicit implementation of open and transparent science principles as assessed by the TOP Factor. We therefore encourage journals to make their requirements more explicit. Furthermore, to promote the recent developments, journals should provide incentives for following open science practices and not only encourage, but make adherence mandatory.

## Supporting information

Supplementary Information S1: Scoring sheet (final)

Supplementary S2: Scoring sheet (intermediate)

Supplementary Information 3: Scoring rubric

## Acknowledgements

This study was supported by the Wellcome Trust (Sir Henry Wellcome Fellowship to M.S.; Wellcome Trust 204686/Z/16/Z), Linacre College, University of Oxford (Junior Research Fellowship to M.S.), by an Erwin-Schroedinger Fellowship of the Austrian Science Fund (FWF; J-4243 to C.B.) and funds from the Freiwillige Akademische Gesellschaft (FAG; C.B.), the Novartis Foundation for Biological-Medical Research (C.B.), and the Psychiatric Hospital of the University of Basel (UPK; C.B.).

## Conflicts of Interest

C.B. is a guest editor for a special issue in *Clocks & Sleep*. M.S. is member of the Editorial Board of *Clocks & Sleep*, and Guest Associate Editor for a Research Topic in *Frontiers in Neurology/Frontiers in Psychiatry*.

**Table 1:**
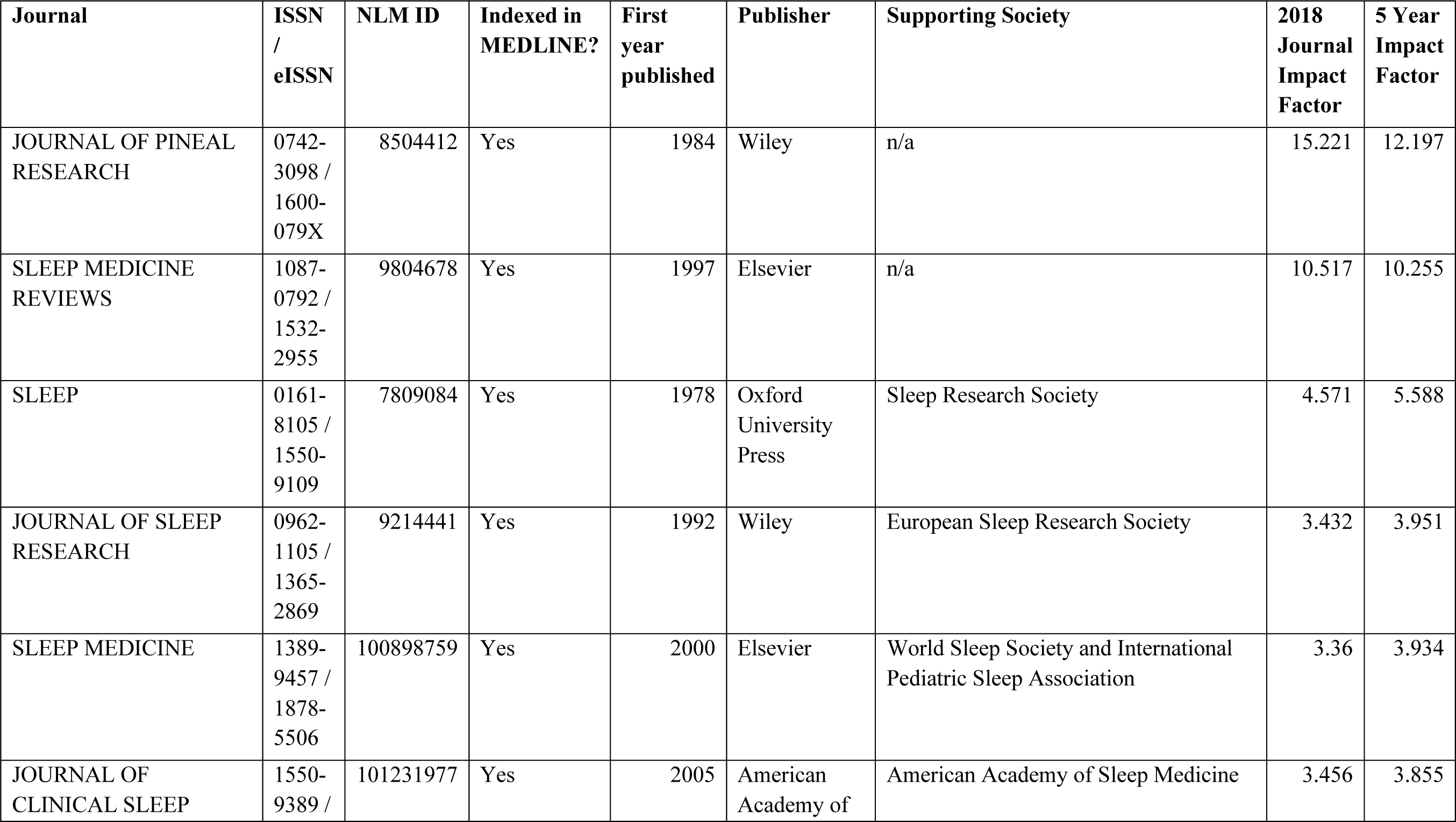
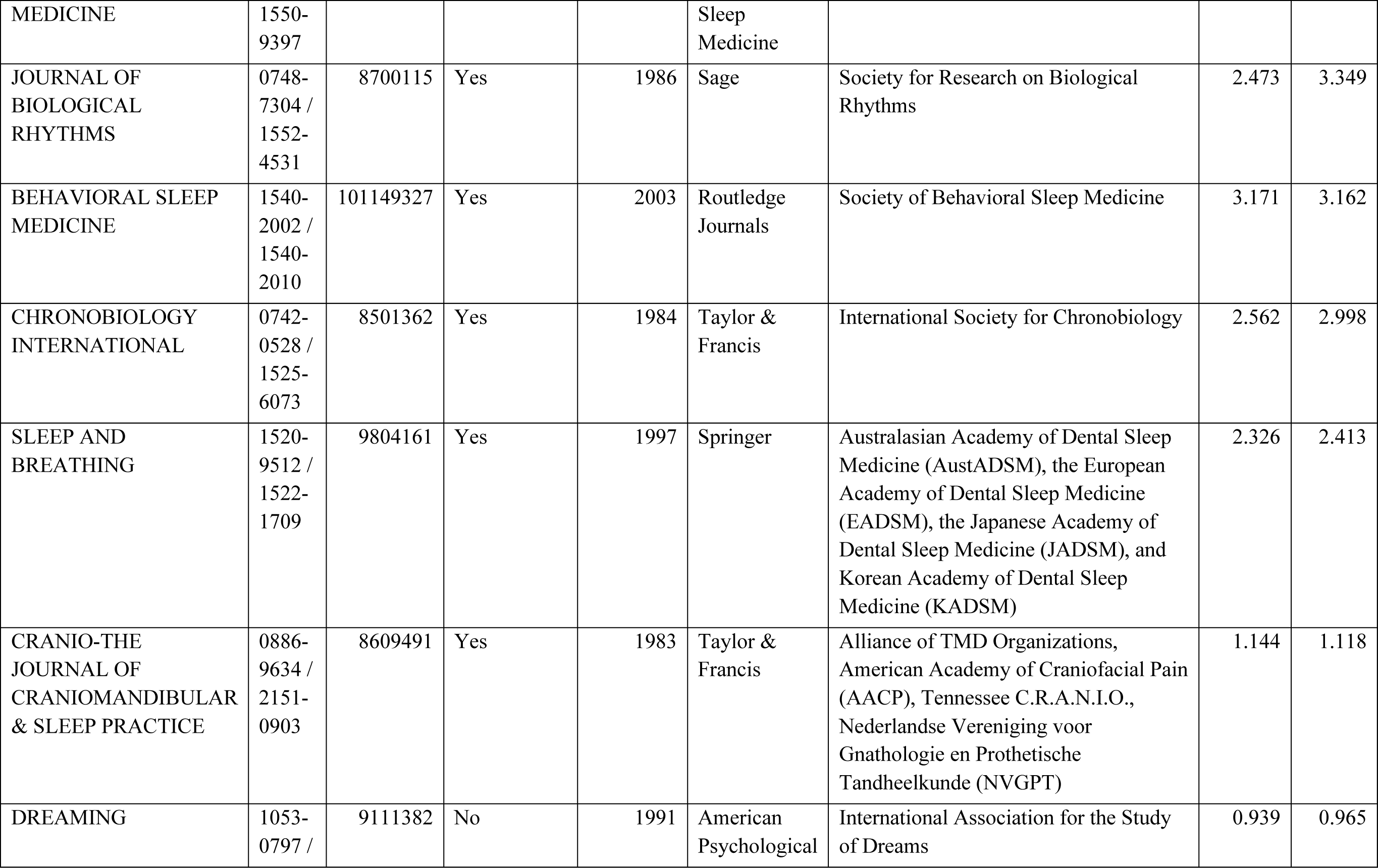
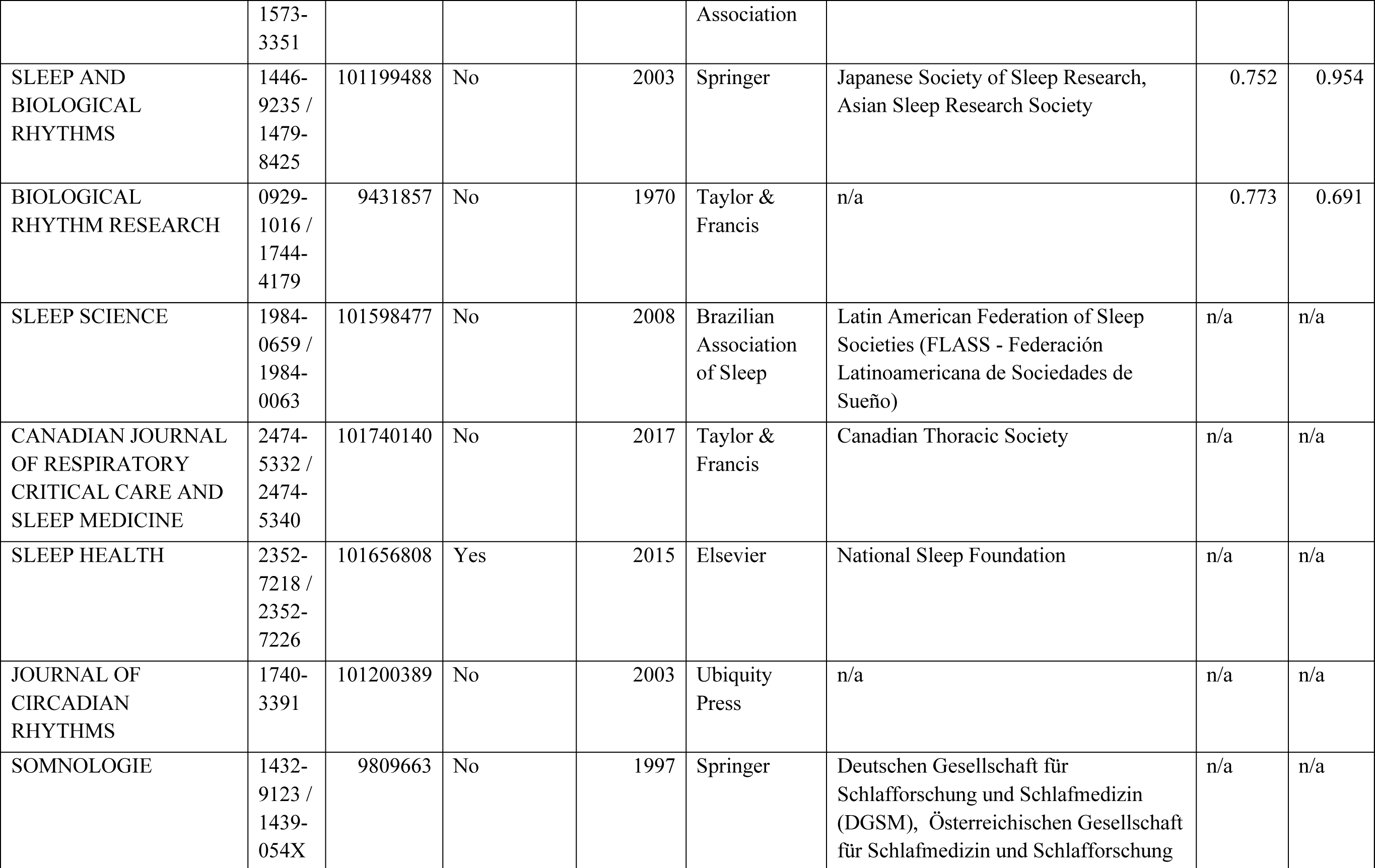
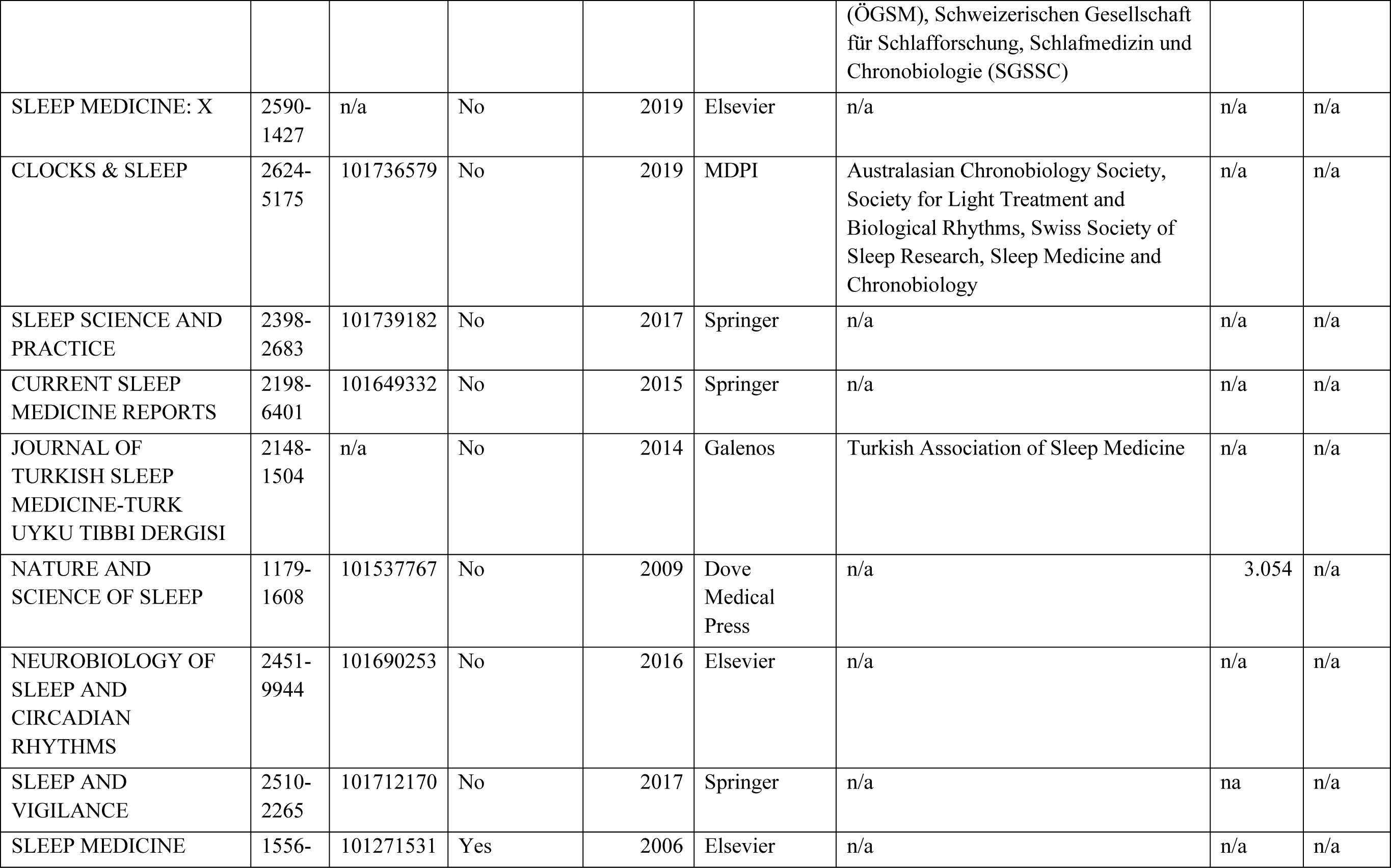
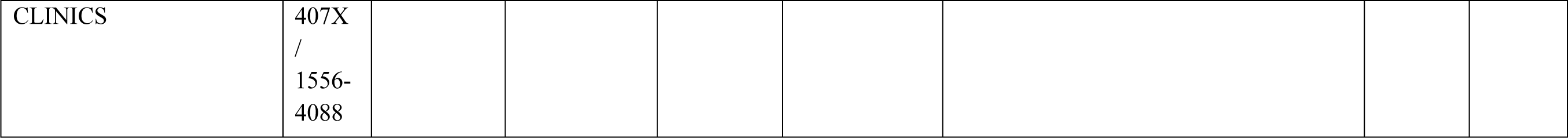
Overview of included journals, sorted by 5-year impact factor. Data included are up to date as of May 2020.

**Table 2:** TOP ratings of included journals, sorted alphabetically by journal name.

| Journal | TOP signatory status (yes, no) | TOP Factor: Data citation (0, 1, 2, or 3) | TOP Factor: Data transparency (0, 1, 2, or 3) | TOP Factor: Analytical code transparency (0, 1, 2, or 3) | TOP Factor: Materials Transparency (0, 1, 2, or 3) | TOP Factor: Reporting guidelines (0, 1, 2, or 3) | TOP Factor: Study prereg (0, 1, 2, or 3) | TOP Factor: Analysis prereg (0, 1, 2, or 3) | TOP Factor: Replication (0, 1, 2, or 3) | TOP Factor: Publication Bias (0, 1, 2, or 3) | TOP Factor: Open science badges (0, 1, or 2) | Total TOP |
| --- | --- | --- | --- | --- | --- | --- | --- | --- | --- | --- | --- | --- |
| BEHAVIORAL SLEEP MEDICINE | No | 2 | 1 | 0 | 0 | 0 | 0 | 0 | 0 | 0 | 0 | 3 |
| BIOLOGICAL RHYTHM RESEARCH | No | 2 | 1 | 0 | 0 | 0 | 1 | 0 | 0 | 0 | 0 | 4 |
| CANADIAN JOURNAL OF RESPIRATORY CRITICAL CARE AND SLEEP MEDICINE | No | 2 | 1 | 0 | 0 | 0 | 1 | 0 | 0 | 0 | 0 | 4 |
| CHRONOBIOLOGY INTERNATIONAL | No | 2 | 1 | 0 | 0 | 0 | 1 | 0 | 0 | 0 | 0 | 4 |
| CLOCKS & SLEEP | No | 2 | 2 | 0 | 2 | 1 | 1 | 1 | 0 | 0 | 0 | 9 |
| CRANIO-THE JOURNAL OF CRANIOMANDIBULAR & SLEEP PRACTICE | No | 2 | 1 | 0 | 0 | 0 | 1 | 0 | 0 | 0 | 0 | 4 |
| CURRENT SLEEP MEDICINE REPORTS | Yes | 0 | 1 | 1 | 1 | 0 | 1 | 0 | 0 | 0 | 0 | 4 |
| DREAMING | No | 0 | 0 | 0 | 0 | 0 | 0 | 0 | 0 | 0 | 0 | 0 |
| JOURNAL OF BIOLOGICAL RHYTHMS | No | 0 | 0 | 0 | 0 | 1 | 1 | 0 | 0 | 0 | 0 | 2 |
| JOURNAL OF CIRCADIAN RHYTHMS | Yes | 0 | 0 | 0 | 0 | 0 | 0 | 0 | 0 | 0 | 0 | 0 |
| JOURNAL OF CLINICAL SLEEP MEDICINE | No | 0 | 0 | 0 | 0 | 0 | 1 | 0 | 0 | 0 | 0 | 1 |
| JOURNAL OF PINEAL RESEARCH | No | 0 | 0 | 0 | 0 | 1 | 1 | 0 | 0 | 0 | 0 | 2 |
| JOURNAL OF SLEEP RESEARCH | No | 0 | 0 | 0 | 0 | 0 | 0 | 0 | 0 | 0 | 0 | 0 |
| JOURNAL OF TURKISH SLEEP MEDICINE-TURK UYKU TIBBI DERGISI | No | 0 | 0 | 0 | 0 | 2 | 1 | 0 | 0 | 0 | 0 | 3 |
| NATURE AND SCIENCE OF SLEEP | No | 0 | 0 | 0 | 0 | 1 | 1 | 0 | 0 | 0 | 0 | 2 |
| NEUROBIOLOGY OF SLEEP AND CIRCADIAN RHYTHMS | Yes | 1 | 0 | 0 | 0 | 1 | 1 | 0 | 0 | 0 | 0 | 3 |
| SLEEP | No | 0 | 0 | 0 | 0 | 1 | 1 | 0 | 0 | 0 | 0 | 2 |
| SLEEP AND BIOLOGICAL RHYTHMS | Yes | 0 | 0 | 0 | 0 | 0 | 1 | 0 | 0 | 0 | 0 | 1 |
| SLEEP AND BREATHING | Yes | 0 | 1 | 1 | 1 | 1 | 1 | 0 | 0 | 0 | 0 | 5 |
| SLEEP AND VIGILANCE | Yes | 1 | 1 | 1 | 1 | 1 | 1 | 0 | 0 | 0 | 0 | 6 |
| SLEEP HEALTH | Yes | 1 | 0 | 0 | 0 | 0 | 0 | 0 | 0 | 0 | 0 | 1 |
| SLEEP MEDICINE | Yes | 1 | 0 | 0 | 0 | 1 | 1 | 0 | 0 | 0 | 0 | 3 |
| SLEEP MEDICINE CLINICS | No | 0 | 0 | 0 | 0 | 0 | 0 | 0 | 0 | 0 | 0 | 0 |
| SLEEP MEDICINE REVIEWS | No | 0 | 0 | 0 | 0 | 0 | 0 | 0 | 0 | 0 | 0 | 0 |
| SLEEP MEDICINE: X | No | 1 | 0 | 0 | 0 | 1 | 1 | 0 | 0 | 0 | 0 | 3 |
| SLEEP SCIENCE | No | 0 | 0 | 0 | 0 | 0 | 1 | 0 | 0 | 0 | 0 | 1 |
| SLEEP SCIENCE AND PRACTICE | No | 2 | 2 | 0 | 2 | 1 | 0 | 0 | 0 | 0 | 0 | 7 |
| SOMNOLOGIE | No | 0 | 0 | 0 | 0 | 0 | 0 | 0 | 0 | 0 | 0 | 0 |
| *Median* |  | 0 | 0 | 0 | 0 | 0 | 1 | 0 | 0 | 0 | 0 | 2.5 |
| *Q1* |  | 0 | 0 | 0 | 0 | 0 | 0 | 0 | 0 | 0 | 0 | 1 |
| *Q3* |  | 1.25 | 1 | 0 | 0 | 1 | 1 | 0 | 0 | 0 | 0 | 4 |
| *IQR* |  | 1.25 | 1 | 0 | 0 | 1 | 1 | 0 | 0 | 0 | 0 | 3 |
| *Minimum* |  | 0 | 0 | 0 | 0 | 0 | 0 | 0 | 0 | 0 | 0 | 0 |
| *Maximum* |  | 2 | 2 | 1 | 2 | 2 | 1 | 1 | 0 | 0 | 0 | 9 |

