## Supplementary Information 3: Scoring rubric for "Transparency and open science reporting guidelines in sleep research and chronobiology journals"

| | 0 | 1 | 2 | 3 |
|---|---|---|---|---|
| **Data citation** | No mention of data citation. | Journal describes citation of data in guidelines to authors with clear rules and examples. | Article requires appropriate citation for data and materials used consistent with the journal's author guidelines. | Article is not published until providing appropriate citation for data and materials following journal's author guidelines |
| **Data transparency** | Data sharing is encouraged, or not mentioned | Articles must state whether or not data are available. Requiring a data availability statement satisfies this level | Articles must have publicly available data, or an explanation why ethical or legal constraints prevent it. | Articles must have publicly available data and must be used to computationally reproduce or confirm results prior to publication |
| **Analytical code transparency** | Code sharing is encouraged, or not mentioned | Articles must state whether or not code is available. Requiring a code availability statement satisfies this level | Articles must have publicly available code, or an explanation why ethical or legal constraints prevent it. | Articles must have publicly available code and must be used to computationally reproduce or confirm results prior to publication |
| **Materials transparency** | Materials sharing is encouraged, or not mentioned | Articles must state whether or not materials are available. Requiring a materials availability statement satisfies this level | Articles must have publicly available materials, or an explanation why ethical or legal constraints prevent it. | Articles must have publicly available materials and must be used to computationally reproduce or confirm results prior to publication |
| **Reporting guidelines** | No mention of reporting guidelines. | Journal articulates design transparency standards. | Journal requires adherence to design transparency standards for review and publication. | Journal requires and enforces adherence to design transparency standards for review and publication. |
| **Study prereg** | Journal says nothing | Articles will state if work was preregistered. | Article states whether work was preregistered and, if so, journal verifies adherence to preregistered plan. | Journal requires that confirmatory or inferential research must be preregistered. |
| **Analysis prereg** | Journal says nothing | Articles will state if work was preregistered with an analysis plan. | Article states whether work was preregistered with an analysis plan and, if so, journal verifies adherence to preregistered plan. | Journal requires that confirmatory or inferential research must be preregistered with an analysis plan. |
| **Replication** | Journal says nothing | Journal encourages submission of replication studies. | Journal will review replication studies blinded to results. | Registered Reports for replications as a regular submission option. |
| **Publication Bias** | Journal says nothing | Journal states that significance or novelty are not a criteria for publication decisions. | Journal will review studies blinded to results. | Journal accepts Registered Reports for novel studies as a regular submission option. |
| **Open science badges** | | Journal awards 1 or 2 open science badges | Journal awards all 3 open science badges | |
